# Kölliker-Fuse/parabrachial complex PACAP - glutamate pathway to the extended amygdala couples rapid autonomic and delayed endocrine responses to acute hypotension

**DOI:** 10.1101/2025.10.16.682741

**Authors:** Vito S. Hernández, Pedro D. Segura-Chama, Limei Zhang

## Abstract

The calyx of Held is a giant axo-somatic synapse classically confined to the auditory brainstem. We recently identified morphologically similar calyx-like terminals in the extended amygdala (EA) that arise from the ventrolateral parabrachial complex and co-express PACAP, CGRP, VAChT, VGluT1, and VGluT2, targeting PKCδ+/GluD1+ EA neurons. Here we asked whether this parabrachial–EA pathway participates in compensation during acute hypotension. In rats given hydralazine (10 mg/kg*, i*.p.), we quantified Fos protein during an early phase (60 min) and a late phase (120 min). Early after hypotension, Fos surged in a discrete subpopulation of the parabrachial Kölliker–Fuse (KF) region and in the EA, whereas magnocellular neurons of the supraoptic and paraventricular nuclei (SON/PVN) remained largely silent. By 120 min, magnocellular SON/PVN neurons were robustly Fos-positive. Confocal immunohistochemistry showed that most Fos+ PKCδ/GluD1 EA neurons were encircled by PACAP+ perisomatic terminals (80.8%), of which the majority co-expressed VGluT1 (88.1%). RNAscope in situ hybridization further identified a selective KF population co-expressing Adcyap1 (PACAP) and Slc17a7 (VGluT1) that became Fos-positive during the early phase. Together these data suggest that a KF^PACAP/VGluT1^ projection forms calyceal terminals around PKCδ/GluD1 EA neurons, providing a high-fidelity route for rapid autonomic rebound to falling blood pressure, while slower endocrine support is subsequently recruited via neurohormone-magnocellular activation. This work links multimodal parabrachial output to temporally layered autonomic-neuroendocrine control.

**Short abstract:** Acute hypotension triggers rapid autonomic compensation followed by slower endocrine support. We identify a Kölliker-Fuse (KF) PACAP/VGluT1 pathway to the extended amygdala (EA) that initiates the fast limb. In hydralazine-treated rats, Fos rose at 60 min in KF and EA but not in magnocellular SON/PVN; by 120 min SON/PVN were strongly Fos-positive. Confocal microscopy showed that ∼81% of Fos+ PKCδ/GluD1 EA neurons were surrounded by PACAP+ calyceal terminals, ∼88% of which co-expressed VGluT1. RNAscope revealed a selective KF Adcyap1/Slc17a7 population that became Fos-positive early. We conclude that KF PACAP/VGluT1 calyces onto PKCδ/GluD1 EA neurons provide a high-fidelity autonomic pathway that precedes vasopressin-mediated endocrine compensation.

## Introduction

The extended amygdala (EA), comprising the capsular division of the central amygdala (CeC) and the oval nucleus of the bed nucleus of the stria terminalis (BSTov), is a hub for rapid viscerosensory integration and for coordinating autonomic, behavioral, and emotional responses to threats and homeostatic challenges (Goehler 2011, Crestani, Alves et al. 2013). These nuclei receive dense inputs from brainstem centers that relay visceral and nociceptive signals – most notably the parabrachial nucleus (PBN) – which alert the EA to internal bodily states and imminent threats (Saper and Loewy 1980, Schwaber, Sternini et al. 1988, Bernard, Alden et al. 1993, Takeuchi, Xie et al. 2004, Jaramillo, Brown et al. 2021). In turn, the CeC and BSTov project to hypothalamic and medullary effectors to orchestrate defensive and stress-related responses; for example, the central amygdala modulates freezing, startle, and autonomic adjustments, including heart-rate and blood-pressure changes, via projections to the rostral ventrolateral medulla (RVLM) and hypothalamus (Holstege, Meiners et al. 1985, Roder and Ciriello 1993, Cobos, Lima et al. 2003, Saha, Drinkhill et al. 2005, Radke 2009). Rapid excitation of CeC neurons, which disinhibit downstream sympatho-excitatory pathways, is therefore well positioned to influence blood-pressure control.

We recently identified a multimodal calyceal synapse in the EA (Zhang, Hernández et al. 2025)formed by PACAP-expressing neurons of the Kölliker-Fuse (KF) nucleus, a ventrolateral subdivision of the parabrachial complex (Eiden 2021, Zhang, Hernandez et al. 2021). These giant axo-somatic terminals co-package glutamate, acetylcholine, and multiple neuropeptides, enveloping PKCδ⁺/GluD1⁺ neurons in the CeC and BSTov. Confocal microscopy, focused-ion-beam scanning electron microscopy (FIB-SEM), and 3-D reconstructions confirmed multiple synaptic specializations onto this PKCδ⁺ subpopulation. The terminals co-express VGluT1, VGluT2, VAChT, PACAP, CGRP, neurotensin, and calretinin, whereas the postsynaptic targets express GluD1, a synaptic adhesion molecule implicated in stabilizing large glutamatergic contacts (Dai, Patzke et al. 2021, Liakath-Ali, Polepalli et al. 2022), suggesting convergence of excitatory, cholinergic, and peptidergic signaling at a single high-fidelity axo-somatic interface.

Located near the pontine–mesencephalic junction, the KF has long been recognized as part of the pontine respiratory (“pneumotaxic”) network that shapes respiratory rhythm and phase transitions and contributes to upper-airway and autonomic control (Varga et al., 2021; Toor et al., 2024; Geerling et al., 2017). Its PACAP– and glutamate-expressing neurons position the KF to integrate sensory signals with autonomic output (Eiden 2021, Zhang, Hernandez et al. 2021).

Here, using a hydralazine-induced acute hypotension model in rats, we show that the KF to CeC/BSTov calyceal pathway is rapidly recruited during the early phase of blood pressure recovery, independently of hypothalamic magnocellular activation. We propose a two-tiered compensation: a fast, high-fidelity KF to EA drive that reinstates sympathetic tone, followed by slower hypothalamo–neurohypophysial support that sustains blood pressure stabilization.

## Results and discussion

Pioneering work had hinted that parabrachial complex to CeC pathway is recruited during acute hypotension (Takeuchi, Xie et al. 2004). To assess the physiological implications of the functional recruitment of KF to EA pathway, we used the hydralazine (HDZ)-induced acute hypotension (Graham, Hoffman et al. 1995) for probing this pathway’s involvement in blood pressure restoration.

The systemic administration of hydralazine (HDZ; 10 mg/kg, i.p. single injection) produced a quick and robust fall in arterial blood pressure (ABP) during the first 60 mins. Averaged mean arterial pressure (MAP) recordings obtained using the CODA HT noninvasive BP system (saline vs HDZ) are shown in the figure 1. The mean ABP fell from 112.8 ± 1.7mmHg to 82.8 ± 3.9 mmHg within the first 5 mins and continued to 68.9 ± 6.4mmHg at the 10 min time point. The mean ABP restored gradually with significant difference disappeared at 90 min time-point after HDZ injection (98.2 ± 6.9 mmHg).

**Figure 1.**
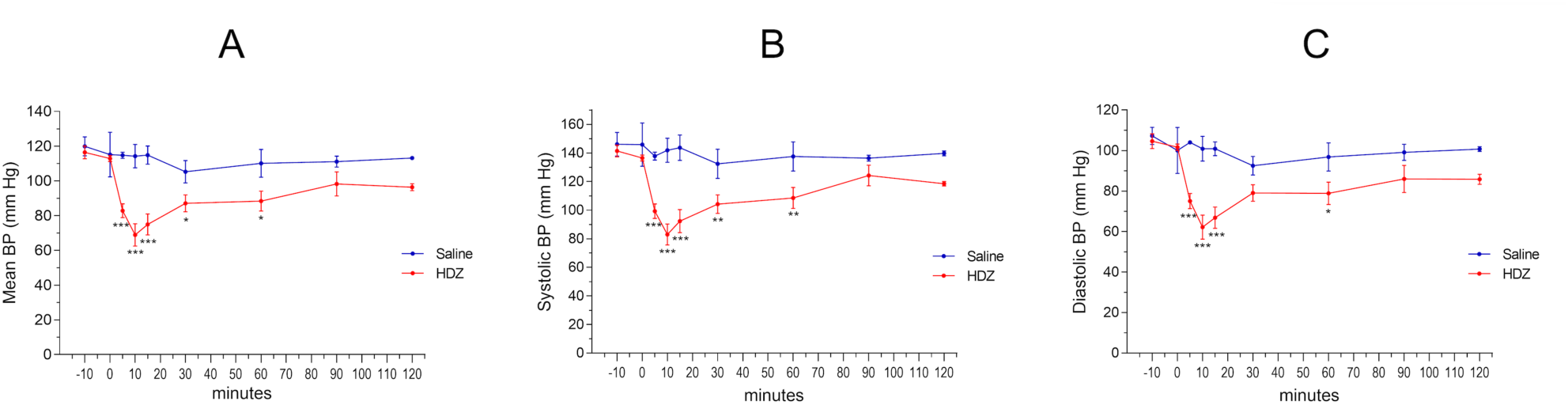
Blood pressure dynamic change as a function of one systemic (intraperitoneal i.p.) hydralazine administration. A. Time course of mean blood pressure (BP) changes measured between 10 min before the i.p. injection (t = 0) to 120 min after the injection. B and C, time courses of systolic and diastolic blood pressure changes respectively (N = 6, n=3). * P < 0.05; ** P < 0.01; ***P < 0.001. Two-way ANOVA followed by Fisher’s LSD multiple comparisons test.

With the acute HDZ hypotensive effect validated, we sought to identify the brain regions engaged during the compensatory response to hypotension, we mapped Fos immunoreactivity at 60 min (“early-Fos”) and 120 min (“late-Fos”) after HDZ injection (Fig. 2A). Perfused-fixed brain at 60 min after the i.p. injection, reflecting RNA fos induction at the beginning the HDZ challenge (Sagar, Sharp et al. 1988, Morgan and Curran 1989, Herrera and Robertson 1996), a distinct subpopulation within the ventrolateral parabrachial complex, within the Kölliker-Fuse nucleus (KF), showed robust Fos induction, together with strong labeling in the capsular central amygdala (CeC) and oval BNST (BSTov) (Fig. 2B3–D3). In contrast, magnocellular neurons of the supraoptic (SON) and paraventricular (PVN) nuclei remained largely quiescent at this time point (Fig. 2E3). By 120 min, however, the SON and PVN exhibited dense nuclear Fos expression (Fig. 2E4), marking a late phase of activation. Quantitative analyses confirmed a significant increase in Fos-positive cell density in KF, CeC, and BSTov at 60 min time point, followed by pronounced labeling in PVN/SON at 120 min (Fig. 2F; one-way ANOVA, different letters indicate p < 0.05). These results delineate a two-stage temporal pattern: an early activation of the KF to EA network that precedes a later recruitment of hypothalamo-neurohypophysial neurons responsible for neurohormone release.

**Figure 2.**
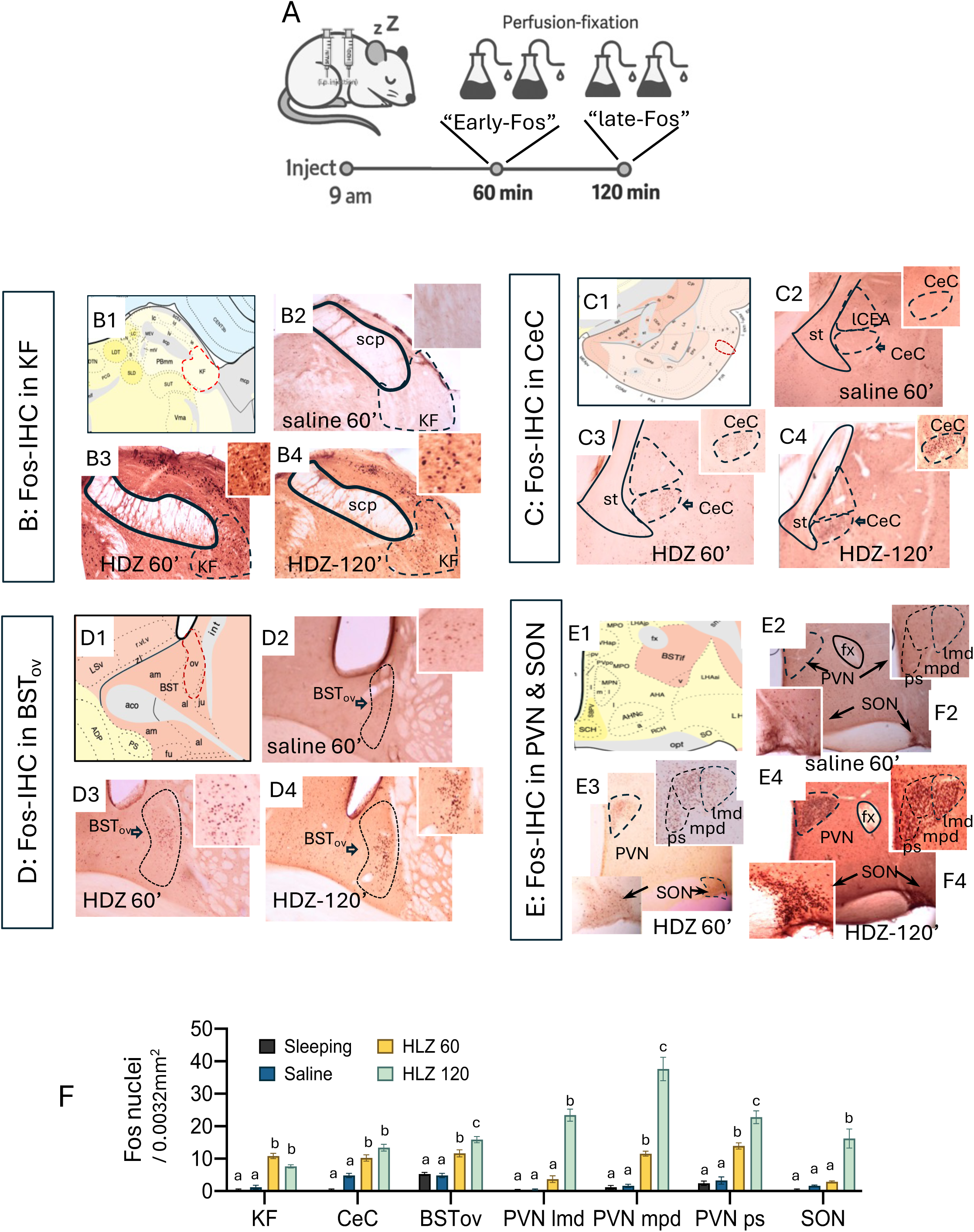
Sequential Fos activation in pontine KF, extended amygdala, and hypothalamic PVN/SON during recovery from acute hypotension. A, Experimental design. Rats were assigned to four groups and perfused at the indicated times after a 9:00 i.p. injection: (i) Sleeping baseline (pentobarbital anesthesia followed by immediate perfusion), (ii) Saline (0.9% NaCl, 1 ml/kg) perfused/fixed at 60 min, (iii) HDZ 60′ (hydralazine, 10 mg/kg in 1 ml/kg saline), perfused/fixed at 60 min, and (iv) HDZ 120′ (hydralazine, 10 mg/kg in 1 ml/kg saline) perfused/fixed at 120 min. B–E, Fos immunohistochemistry (IHC) across regions. For each block, panel 1 shows the corresponding <u>Swanson atlas level, 2004</u>; panel 2 shows Saline 60′; panel 3 shows HDZ 60′; panel 4 shows HDZ 120′. B, KF (Kölliker–Fuse nucleus, ventrolateral parabrachial complex). C, CeC (capsular central amygdala). D, BSTov (oval nucleus of the bed nucleus of the stria terminalis). E, PVN & SON (paraventricular and supraoptic nuclei of the hypothalamus; PVN subnuclei labeled using Swanson nomenclature). Insets illustrate nuclear Fos at higher magnification. F, Quantification. Fos-positive nuclei (mean ± SEM) are plotted as cells per 0.0032 mm² (60 µm × 53 µm field) for each region and group (Sleeping, Saline 60′, HDZ 60′, HDZ 120′). Different letters above bars indicate significant differences (two-way ANOVA within region followed by Tukey post hoc, α = 0.05). See Methods for sample sizes and counting criteria.

To determine the cellular targets activated during the early phase, we combined Fos immunofluorescence with markers of the calyceal input previously described in the extended amygdala. At 60 min post-HDZ, numerous Fos-positive nuclei were observed within the CeC, surrounded by ring-like PACAP-immunoreactive terminals (Fig. 3A). High-resolution confocal imaging revealed that these perisomatic rings correspond to VGluT1⁺/PACAP⁺ axo-somatic structures apposed to PKCδ⁺ neurons (Fig. 3B). Many of these terminals also contained CGRP and formed contacts adjacent to GluD1-positive postsynaptic domains (Fig. 3C,D). Quantitative analysis showed that 80.8 % of Fos-positive neurons were surrounded by PACAP⁺ calyceal terminals and 88.1 % of these perisomatic rings co-expressed VGluT1 (Fig. 3E,F). These data demonstrate that the PACAP/VGluT1 calyceal input to PKCδ/GluD1 CeC neurons is selectively engaged during the early compensatory phase of acute hypotension.

**Figure 3.**
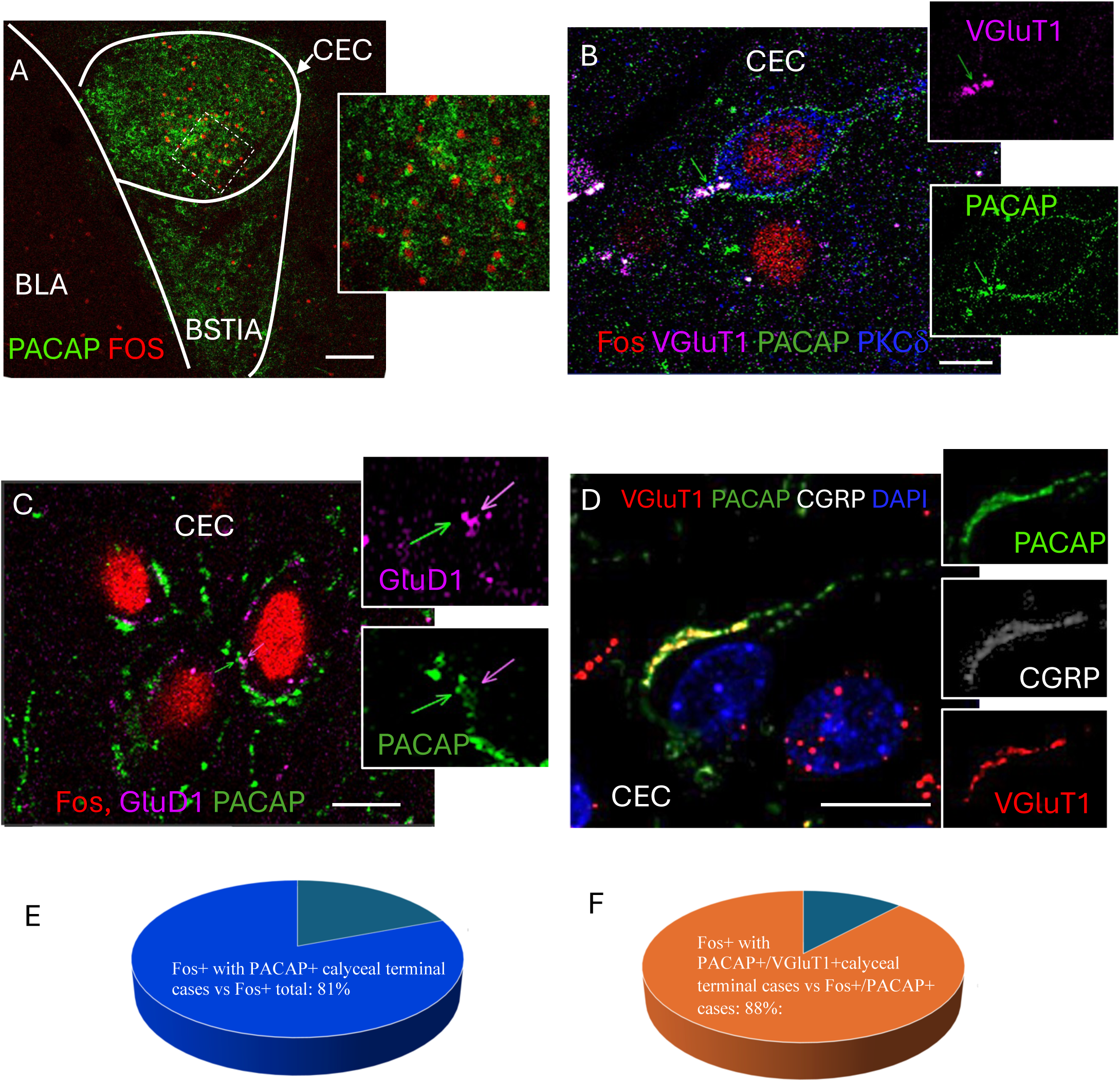
Early-phase Fos reveals selective engagement of PACAP/VGluT1 calyceal inputs onto PKCδ/GluD1 neurons in the CeC. A, Low magnification. Fos-positive nuclei (red) in the capsular central amygdala (CeC) 60 min after hydralazine. Inset: Fos+ nuclei (red) encircled by ring-like PACAP-immunoreactive (PACAP-ir) perisomatic terminals (green). B–D, High-resolution confocal images. B. A PKCδ-immunoreactive (PKCδ-ir; blue) neuron expressing Fos is surrounded by PACAP/VGluT1 co-expressing calyceal terminals (green/magenta). C. PACAP-ir axonal rings are apposed by GluD1-ir puncta on the postsynaptic soma (arrows), consistent with a calyceal axo-somatic contact. D. Calyceal terminals co-express PACAP, CGRP, and VGluT1 (right panels show single channels). E–F, Quantification. E. Proportion of Fos+ CeC neurons encircled by PACAP+ calyceal terminals: 80.8% (42/52 cells). F. Among Fos+/PACAP+ cases, proportion with VGluT1^+ calyceal co-labeling: 88.1% (37/42 cells).Scale bars: A, 100 µm; B–D, 10 µm. Confocal images acquired at 1 Airy unit (z-step as in Methods). Abbreviations: BLA, basolateral amygdala; BSTIA, bed nucleus of the stria terminalis, intra-amygdaloid component; DAPI, nuclear counterstain.

To identify the upstream neurons giving rise to the PACAP/VGluT1 calyceal terminals, we examined gene expression in the KF using single– and duplex-RNAscope in situ hybridization. Adcyap1 (PACAP mRNA)-positive neurons formed a dense cluster within the ventrolateral parabrachial complex (Fig. 4A), and many also expressed Slc17a7 (VGluT1 mRNA; Fig. 4B,E). Duplex assays combining Fos with Adcyap1 or Slc17a7 revealed that a subpopulation of PACAP⁺/VGluT1⁺ KF neurons became Fos-positive at 60 min following HDZ (Fig. 4C,D). Additional duplex reactions confirmed partial co-expression of Adcyap1 and Calca (CGRP) (Fig. 4F), consistent with the mixed neurochemical profile of the calyceal terminals in the CeC. Together, these data identify a discrete PACAP/VGluT1-expressing KF subpopulation that is activated during the early phase of hypotensive challenge and likely constitutes the source of the PACAPergic calyceal projection to the extended amygdala.

**Figure 4.**
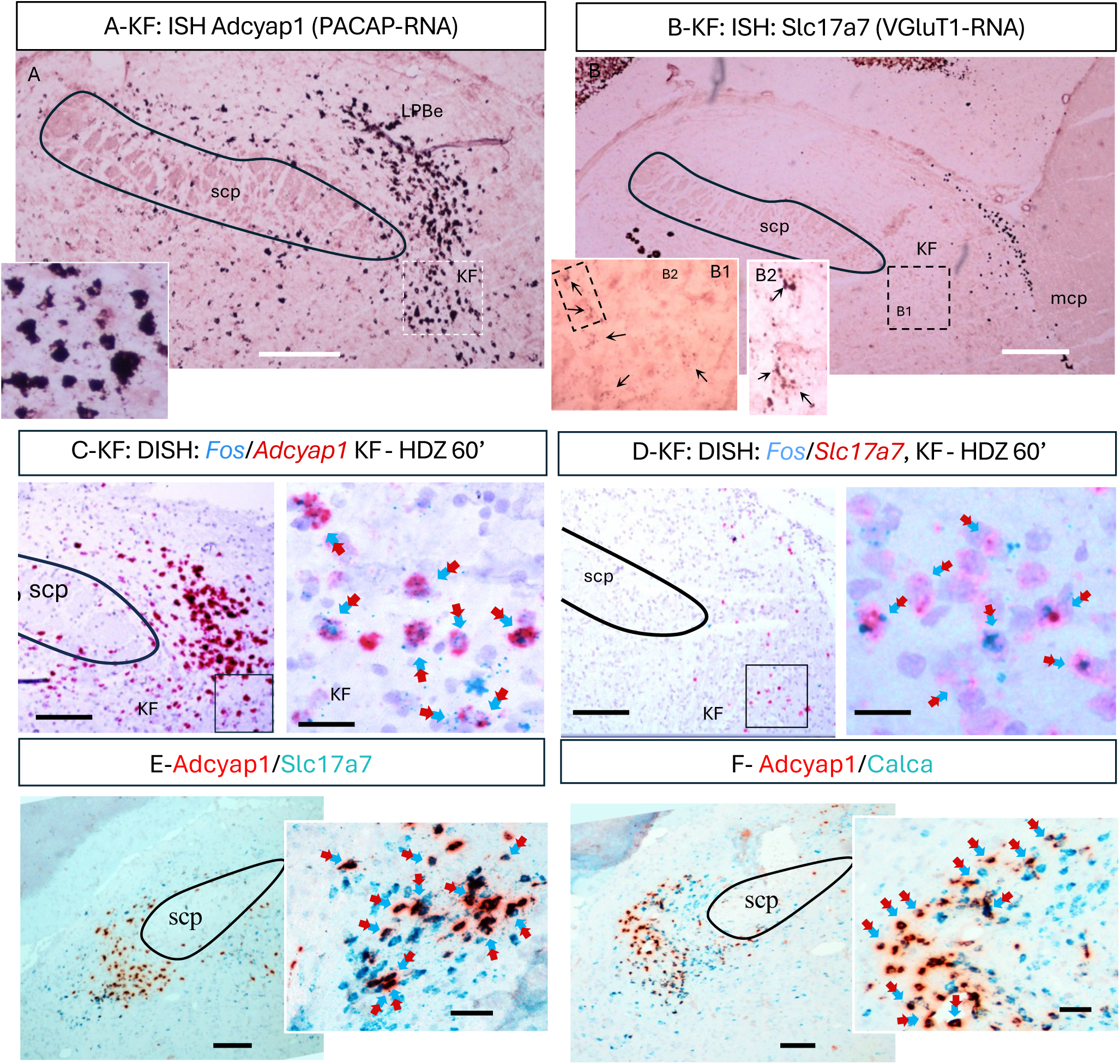
A subpopulation in the pontine Kölliker-Fuse nuclei (KF) which expressed mRNA of PACAP/VGluT1 engaged in the early phase of the acute hypotension event. A and B: ISH using high sensitivity/low background single cell labeling with RNAscope brown-kit method revealed a peculiar cell population of rat brain located in the ventral subdivision of the pontine parabrachial complex, at bregma –9.12mm to – 9.24mm, medio-lateral 2.90mm – 3.00mm and dorso-ventral 7.40mm to 7.60mm, between the superior and medial cerebellar peduncles (scp and mcp, respectively), with large cell bodies, expressing Adcyap1 (mRNA for PACAP, panel A and inset) and Slc17a7 (mRNAs for VGluT1, panel B and inset, where the Slc17a7 positive grains can be clearly seen on and surround the Nissl stained nuclei, white arrows). C and D: with RNAscope duplex method (DISH) it is evident that this KF^PACAP/VGluT1^ population is fos positive at the early raising phase. E and F, DISH reactions show the co-expression of Adcyap1 (PACAP) co-expressing Slc17a7 (VGluT1) and Calca (CGRP). Scale bars: A and B, 200µm, C-F: left panels, 50µm, right panels: 25µm. Single KF neuron projections links pontine source to EA calyceal targets. In vivo juxtacellular labeling of a PACAP-positive neuron in the Kölliker–Fuse (KF) revealed three long-range axonal branches: (i) a caudo-medial branch toward the medulla, (ii) a dorsal branch within the pons, and (iii) a rostral branch that coursed through major conduits: superior cerebellar peduncle (scp), to dorsal tegmental bundle entering the capsular central amygdala (CeC), and continued via the ansa peduncularis to the BSTov. Along this route we observed labeled fibers within the scp, ring-like (“calyceal-like”) perisomatic terminals in BSTov, and neurobiotin/PACAP-apposed terminals in CeC, supporting a KF origin for the PACAP-rich perisomatic inputs identified in Figures 3–4.

**Figure 5.**
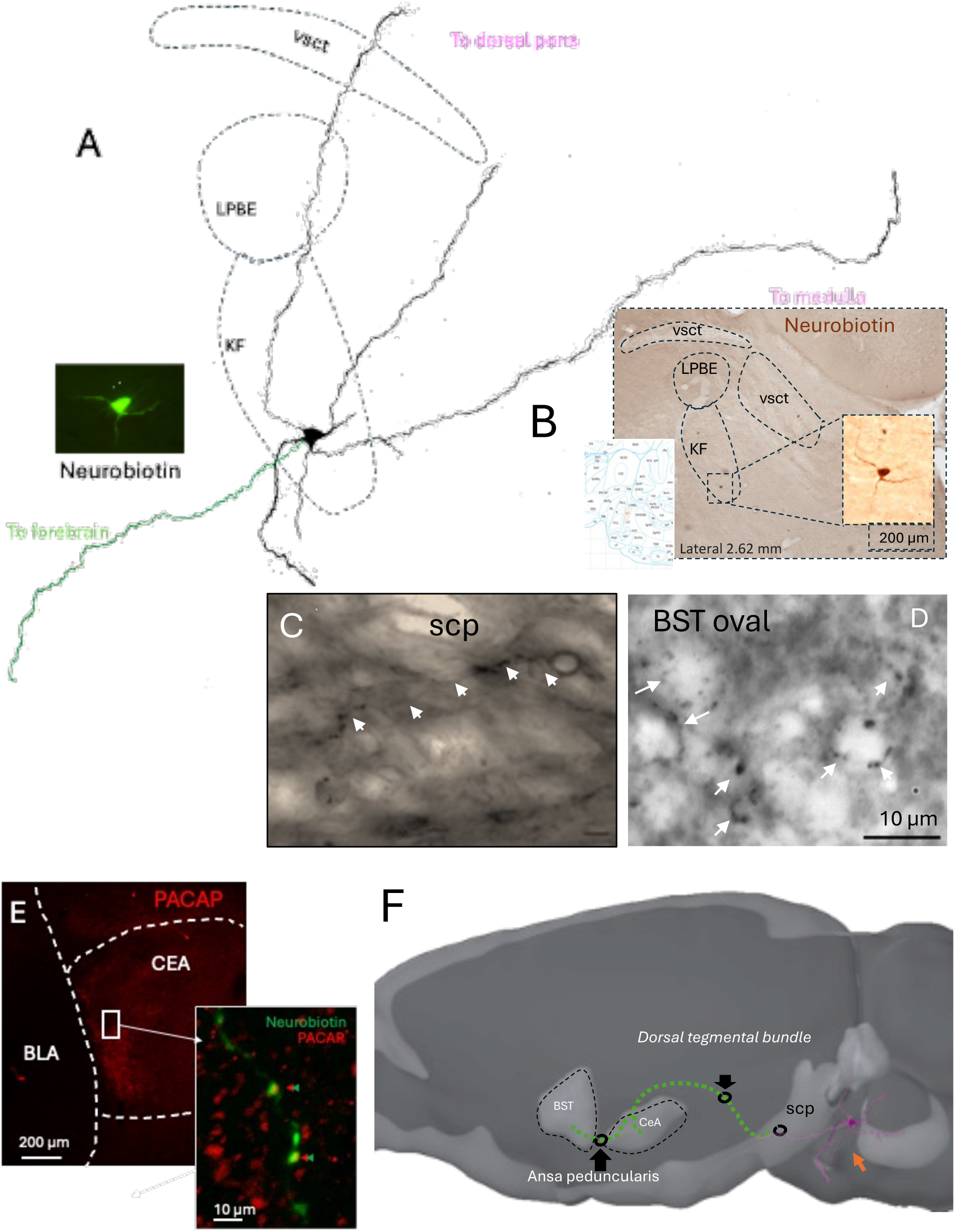
A juxtacellularly labeled KF neuron with long-range projections to medulla, dorsal pons, and forebrain targets. A, Camera lucida reconstruction. Ten serial 60-µm sections were aligned to reconstruct a single Kölliker–Fuse (KF) neuron filled juxtacellularly with neurobiotin. The axon issues three major branches coursing medially/caudally to the medulla, dorsally within the pons, and rostrally toward the forebrain. Local landmarks are outlined (LPBe, external division of the lateral parabrachial nucleus; vsct, ventral spinocerebellar tract). B, Soma location. Bright-field micrograph showing the labeled KF neuron at its origin within the ventrolateral parabrachial complex (inset: higher magnification of the soma). Scale bar, 200 µm. C, Axon within scp. Neurobiotin-labeled axon (arrows) traveling in the superior cerebellar peduncle (scp). Scale bar, 10 µm. D, Ring-like terminal in BSTov. Enlarged neurobiotin-positive perisomatic “calyceal-like” terminal in the oval BNST (BSTov). Scale bar, 10 µm. E, Forebrain terminals co-localize with PACAP. Left: PACAP immunofluorescence (red) in the capsular central amygdala (CeC) and adjacent basolateral amygdala (BLA). Right inset: merged confocal image showing neurobiotin-labeled terminals (green) apposed to PACAP-immunoreactive perisomatic profiles (red) in the CeC. Scale bars: left 200 µm, inset 10 µm. F, Schematic (sagittal). Summary of the reconstructed trajectory: the rostrally directed branch enters major brain conduction systems—scp → dorsal tegmental bundle (DTB) → CeC—and continues via the ansa peduncularis to the BSTov; additional branches descend toward the medulla. This single-cell anatomy supports a KF origin for PACAP-positive calyceal innervation of the extended amygdala. Abbreviations: BLA, basolateral amygdala; CeC, capsular central amygdala; DTB, dorsal tegmental bundle; KF, Kölliker–Fuse nucleus; LPBe, lateral parabrachial nucleus, external division; scp, superior cerebellar peduncle; BSTov, oval nucleus of the bed nucleus of the stria terminalis; vsct, ventral spinocerebellar tract.

## Discussion

Acute hypotension elicits a rapid autonomic rebound followed by slower endocrine support. Here we identify a discrete Kölliker–Fuse (KF) PACAP/VGluT1 pathway to the extended amygdala (EA) that is engaged during the early phase of blood-pressure recovery, preceding hypothalamo– neurohypophysial activation. Our findings reveal a previously unrecognized synaptic mechanism that couples a rapid autonomic response with a delayed neuroendocrine compensation after acute hypotension. The identification of a PACAP/VGluT1/CGRP calyceal projection from the Kölliker–Fuse (KF) nucleus to PKCδ⁺/GluD1⁺ neurons of the extended amygdala (EA) introduce a new element into the organization of central networks that control blood pressure. This discovery expands the classical model of cardiovascular homeostasis beyond medullary baroreflex circuits, incorporating a high-fidelity pontine–amygdaloid route capable of influencing sympathetic outflow with exceptional temporal precision.

The biphasic Fos pattern observed i.e. early activation in the KF and EA followed by delayed recruitment of vasopressinergic magnocellular neurons, suggests a two-stages temporal hierarchy in the brain’s response to falling blood pressure. In the first stage, PACAP/VGluT1/CGRP calyceal input rapidly excites PKCδ⁺ neurons in CeC and BSTov, which are strategically positioned to modulate sympathoexcitatory outputs through disinhibition of hypothalamic and medullary targets (Crestani, Alves et al. 2013, Son, Filosa et al. 2013). This fast relay may provide the neural substrate for the “autonomic rescue” phase, restoring arterial pressure before endocrine mechanisms become engaged. The functional organization of the CeA microcircuit offers a plausible mechanism by which KF input could translate into sympathoexcitatory drive. PKCδ⁺ neurons in the central capsular subdivision (CeC/CeL) provide inhibitory control over output neurons in the medial CeA (CeM) which themselves project to the rostral ventrolateral medulla (RVLM) and other premotor autonomic nuclei (Cobos, Lima et al. 2003, Bou Farah, Bowman et al. 2016). Activation of PKCδ⁺ neurons suppresses CeM activity, thereby attenuating CeM-mediated inhibition of brainstem targets (Haubensak, Kunwar et al. 2010). In the context of acute hypotension, PACAP/VGluT1 calyceal input from the KF to PKCδ⁺ CeC neurons could therefore reduce CeM output, disinhibiting RVLM sympathetic premotor neurons and facilitating the rapid restoration of arterial pressure. This interpretation reconciles the inhibitory nature of PKCδ⁺ neurons with the net excitatory outcome on sympathetic tone, linking the KF→EA projection anatomically and functionally to the baroreflex recovery phase.

The identification of PKCδ⁺ neurons as principal recipients of KF input is consistent with their well-established role as inhibitory gatekeepers within the central amygdala microcircuitry. Haubensak et al. used optogenetic tools to demonstrate that PKCδ⁺ neurons in the lateral/central amygdala (CeL/CeC) exert potent inhibitory control over medial output neurons (CeM), forming a disinhibitory switch that governs behavioral and autonomic responses to salient stimuli. Within this framework, the KF PACAP/VGluT1 calyces described here could represent a high-fidelity excitatory drive onto these inhibitory gate neurons, biasing the circuit toward rapid activation of downstream sympathoexcitatory pathways when visceral homeostasis is threatened. Such an arrangement would allow the extended amygdala to function as a dynamic relay, integrating pontine visceral inputs with affective context while gating the flow of information to medullary autonomic centers. Thus, our findings extend the principle established by Haubensak and colleagues—of PKCδ⁺ neurons as modulatory nodes—beyond conditioned fear paradigms to the realm of cardiovascular control, highlighting their broader role in orchestrating state-dependent autonomic output.

The second delayed stage, engages magnocellular SON/PVN neurons to ensure sustained vasopressin-mediated volume recovery and allostatic stabilization (Leng and Russell 2019). The delayed recruitment of magnocellular neurons in the PVN and SON likely reflects the operation of local circuits that gate hypothalamic output during stress and autonomic challenges. As proposed by Herman et al. (2002), the PVN functions not as a simple relay but as a microcircuit hub where excitatory and inhibitory inputs from limbic and brainstem sources converge onto GABAergic and glutamatergic interneurons before influencing neurosecretory and preautonomic neurons (Herman, Tasker et al. 2002). Such organization provides a mechanism for temporal filtering, in which rapid autonomic inputs, potentially relayed through the KF – EA pathway activate upstream excitatory drives that are initially counterbalanced by strong local inhibition. Only when this inhibitory tone is lifted, or when excitatory inputs summate beyond a critical threshold, do PVN magnocellular neurons become engaged, producing the late phase of Fos induction and vasopressin release. This ensures that neuroendocrine compensation occurs only after the initial autonomic rebound has stabilized arterial pressure, thereby preventing redundant or excessive activation of vasopressinergic systems.

Converging anatomical evidence reinforces the notion that the extended amygdala exerts direct influence over medullary sympathetic control centers. For instance, anterograde tracing from the rat central amygdala (CeA) combined with with electron microscopy demonstrated a direct CeA→RVLM projection making synaptic contacts onto phenylethanolamine-N-methyltransferase–positive (C1) medullary sympathetic premotor neurons activated by hypotension (Saha et al., 2005). Using cholera toxin beta, CeA afferents to the ventrolateral medulla were confirmed, further supporting a continuous extended-amygdala→VLM/RVLM axis. These studies positioned the CeA as and efferent component of the cardiovascular reflex integration. Also, it has been shown that somatostain containig projections from CeA also reach the RVLM, indicating the existence of a heterogenic amygdalo-medullary pathway that potentially has effects of modulating the sympathetic tone (Cobos, Lima et al. 2003, Bou Farah, Bowman et al. 2016).

Additionally, it has been shown In the cat that the BNST also sends descending fibers to ventrolateral medullary autonomic fields including the lateral tegmental field, nucleus of the solitary tract, and dorsal vagal complex, overlapping ventrolateral medullary regions that flank the RVLM, also the authors discuss the similarity of the terminal fields innervated by the CeC and the BNST, suggesting the possibility to consider both regions as aone anatomical entity (Holstege, Meiners et al. 1985).

The calyx-like terminals described here resemble those of the auditory brainstem in scale and transmitter complexity, but they occur in a forebrain circuit governing visceral / autonomic state rather than sound localization. Their morphology and multi-transmitter composition imply exceptionally reliable transmission with the potential for graded modulation by neuropeptides. PACAP, co-released with glutamate, likely enhances presynaptic Ca²⁺ influx and postsynaptic excitability, thus amplifying the gain of the autonomic response (Zhang, Hernandez et al. 2021, Zhang, Hernández et al. 2025). The co-presence of VAChT in these PACAP/CGRP terminals, suggests synergistic recruitment of peptidergic and cholinergic modulation, conferring both precision and persistence. Functionally, this arrangement may enable the EA to detect visceral perturbations not merely as sensory inputs but as urgent salience signals requiring rapid autonomic adjustment.

The juxtacellular reconstruction of an individual KF neuron provides direct anatomical continuity from the pontine source to forebrain targets: a rostrally directed axon traveling through the scp and dorsal tegmental bundle to reach the CeC and then the BSTov via the ansa peduncularis, with calyceal-like terminals observed in BSTov and PACAP-apposed terminals in CeC. This single-cell evidence dovetails with our RNAscope and confocal findings— PACAP/VGluT1 KF neurons activated early (Fig. 4) and PACAP/VGluT1 calyces encircling Fos+ PKCδ/GluD1 EA neurons (Fig. 3)—thereby closing the loop from molecular identity → activation → pathway anatomy → target engagement. Together with the sequential Fos mapping (Fig. 2), the data support a model in which a high-fidelity KF→EA channel initiates rapid autonomic recovery, while magnocellular SON/PVN neurons provide delayed endocrine stabilization.

By positioning the EA as a recipient of KF autonomic drive, our results bridge two traditionally distinct domains viscerosensory control and emotional processing. The same PKCδ⁺ EA neurons involved in hypotensive compensation also regulate fear and anxiety states (Duvarci, Bauer et al. 2009, Duvarci and Pare 2014, Hevesi, Zelena et al. 2021) Thus, the calyceal pathway could represent an anatomical substrate through which internal bodily challenges engage affective circuits, giving rise to the subjective awareness of “bodily threat” (Critchley, Mathias et al. 2002). The dual role of PACAP in both stress (Stroth, Holighaus et al. 2011, Zhang and Eiden 2019) and cardiovascular regulation (Dalsgaard, Hannibal et al. 2003) supports this idea: PACAP signaling in the amygdala and BNST has been implicated in anxiety and panic disorders (Missig, Mei et al. 2017, Li, Andero et al. 2023). Our data therefore offer a mechanistic framework for understanding how visceral disturbances can trigger emotional arousal, unifying autonomic and affective physiology.

This study introduces the first direct evidence of a multimodal calyceal synapse operating within a limbic–autonomic network. Beyond hypotension, such architecture may contribute to cardiorespiratory coupling during hypoxia (Toor, Burke et al. 2024) or to stress-induced sympathetic activation (Damasceno, Takakura et al. 2015). Future electrophysiological work would be key to determine whether PACAP-dependent facilitation at these terminals shows activity-dependent plasticity, potentially explaining persistent sympathetic bias in chronic stress or hypertension. Likewise, identifying homologous KF to EA pathways in humans (Lavezzi, Ottaviani et al. 2004)may clarify the neural basis of dysautonomic states and anxiety-related cardiovascular symptoms.

### Limitations and future directions

Our study is correlative with respect to causality: we infer the pathway’s contribution from Fos mapping and anatomical convergence. Direct tests will require temporally precise inhibition or activation of KF PACAP/VGluT1 neurons and selective disruption of EA calyces (e.g., synaptotagmin manipulation, targeted peptide receptor blockade). Continuous telemetry would complement tail-cuff measures to capture second-to-second dynamics. Finally, identifying the downstream EA efferents that mediate sympathetic rebound (e.g., projections to RVLM or hypothalamic premotor areas) will clarify the effector arm.

### Conclusion

We define a KF PACAP/VGluT1 projection that forms calyceal terminals onto PKCδ/GluD1 neurons in the extended amygdala and is selectively engaged during the early phase of recovery from acute hypotension. This pathway provides a mechanistic substrate for rapid autonomic compensation that precedes and complements slower vasopressin-dependent endocrine support, revealing a temporally stratified architecture for cardiovascular homeostasis.

## Materials and methods

Adult (>60 days) male Wistar rats were obtained from the vivarium of the School of Medicine, National Autonomous University of Mexico. Rats were housed under controlled temperature and humidity on a 12-h light cycle (lights on at 07:00, off at 19:00) with ad libitum access to standard chow and water. All procedures were approved by the School of Medicine’s Ethics Committee (License: CIEFM-079-2020).

## Blood Pressure Measurements

The blood pressure (BP) measurements in conscious rats were done between 10:00 a.m. and 03:00 p.m. via a non-invasive tail-cuff CODA system (Kent Scientific Corp., USA). Animals were acclimated to restraining device and the sensing tail-cuff during 30 min to avoid unnecessary stress in the course of measurements. During recording, the occlusion cuff and the volume pressure recording (VPR) sensor was positioned on the tail base while the rats were maintained in a heater platform set to 30 °C to induce vasodilation, which enhances the blood flow to the tail. Data of BP (the average of 3 times) were evaluated two times at basal level and 5, 10, 15, 30 and 60, 90 and 120 minutes after hydralazine (10 mg/Kg-1) or saline (0.9 % NaCl, 1 mL/Kg-1) intraperitoneal administration. CODA software displays the systolic, diastolic and mean blood pressure data in mmHg units. Data are expressed as mean ± SEM of BP measurements in mmHg of at least 3 rats per condition and indicated time. Statistical analysis was performed using two-way ANOVA followed by Fisher’s LSD multiple comparisons post hoc test, a statistically significant difference was set with a P value < 0.05.

### Immunohistochemistry

Rodents were perfused via the ascending aorta with 0.9% NaCl followed by 4% paraformaldehyde. Brains were sectioned at 70 µm using a Leica VT1200 vibratome. Sections were washed in phosphate buffer (PB), blocked for 2 hours in TBST (0.05 M Tris, 0.9% NaCl, 0.3% Triton-X) plus 10% normal donkey serum, and incubated overnight at 4°C with primary antibodies (see SI table 1 diluted in TBST with 1% serum. After washing, sections were incubated for 2 hours at room temperature with appropriate biotinylated or fluorescent secondary antibodies (1:500; Vector Laboratories). Brightfield images were acquired using a Nikon Eclipse E600, and confocal images were acquired with a Leica Stellaris microscope.

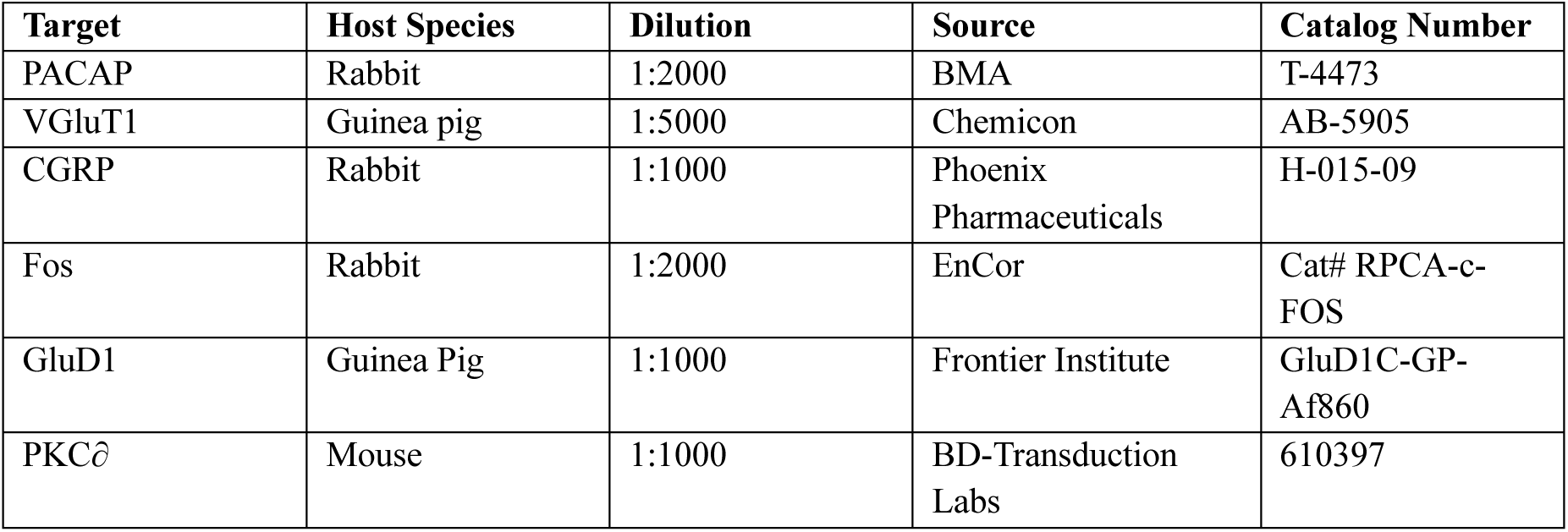
Primary antibodies.

### Fos counting

regions of interest (ROIs) were delineated on bright-field images using Swanson atlas landmarks (KF, CeC, BSTov, PVN subnuclei, SON). Within each ROI, non-overlapping fields of 60 µm × 53 µm (0.0032 mm²) were sampled bilaterally and quantified as cells per field. A neuron was scored Fos⁺ only when it contained a discrete intranuclear DAB deposit fully enclosed by the hematoxylin counterstain, with round/oval morphology and intensity clearly above local background; cytoplasmic staining, diffuse puncta, and glial-like profiles were excluded. Unbiased counting-frame rules were applied (top/left borders inclusive; bottom/right borders exclusive) to avoid edge effects; fields with tissue folds, tears, or vascular artifacts were omitted. For each animal, all fields from all available sections within the defined atlas levels were averaged to yield a single per-animal value for each ROI; these per-animal means were used for statistics. Counting was performed blind to group identity, with identical illumination and threshold settings maintained across groups for a given staining batch.

### Statistics for Fos counts

we analyzed per-animal means using a two-way ANOVA with factors Region (KF, CeC, BSTov, PVN_lmd/ps/mpd_, SON) and Group (Sleeping, Saline 60′, HDZ 60′, HDZ 120′). When the Group main effect or region × group interaction was significant, we probed simple effects within each region by one-way ANOVA followed by Tukey’s HSD pairwise comparisons (α=0.05, two-tailed). In bar graphs, different letters above bars indicate groups that differ significantly within that region. Data are shown as mean ± SEM; the animal is the statistical unit. Normality were checked; no corrections were required.

### RNAscope In Situ Hybridization

rat brains were rapidly dissected, flash-frozen, and stored at −80 °C. Coronal sections (12 µm) were cut at −20 °C and mounted onto slides. RNAscope was performed using manufacturer instructions (Advanced Cell Diagnostics). Sections were fixed, dehydrated, and treated with Protease IV before hybridization with target probes at 40 °C for 2 h. The following probes were used: Adcyap1, Slc17a7, Slc17a6, and Calca. After hybridization, slides were washed, amplified, counterstained with DAPI, and coverslipped. Imaging was performed using a Leica Stellaris confocal microscope with a 20× objective.

### Juxtacellular labeling and histological analysis

For this study, juxtacellular recording and labeling was performed according to previous protocols (Pinault 1994, Hernandez, Vazquez-Juarez et al. 2015, Zhang, Hernandez et al. 2016, Zhang, Hernandez et al. 2018). Juxtacellular recordings were performed in adult male rats under urethane anesthesia (1.5 g/kg, i.p.). Glass micropipettes (tip resistance 25–40 MΩ) were filled with 1.5% Neurobiotin (Vector Laboratories) dissolved in 0.15 M NaCl. Neurons were recorded extracellularly in the Kölliker-Fuse (KF) region of the parabrachial nucleus, identified by stereotaxic coordinates (−9.2mm posterior to bregma, 2.9mm lateral, and 7.5 mm ventral from the brain surface). Following electrophysiological identification, single neurons were labelled by juxtacellular iontophoresis of Neurobiotin using positive current pulses (200 ms, 1–10 nA, 50% duty cycle) applied for 5 minutes, once modulation of the firing rate was observed.

After a survival period of 6 hours, animal was deeply anesthetized and perfused transcardially with 0.9% NaCl followed by 4% paraformaldehyde in PBS + 15% v/v of a saturated solution of picric acid. Brains were sectioned sagittally at 70 µm using a vibrating microtome. Free-floating sections were processed first with Streptavidin coupled to Alexa 488, and then turned into a permanent DAB labelling using an avidin-biotin–peroxidase complex (ABC Elite, Vector Laboratories) and 3,3′-diaminobenzidine (DAB) as chromogen. To reveal co-localization of the labelled axons with pituitary adenylate cyclase-activating polypeptide (PACAP), some sections with labeled axonal profiles within the amygdala were incubated with Streptavidin-488 together with a rabbit anti-PACAP antibody (1:1000; BMA, cat. T-4473) followed by a fluorophore-conjugated secondary antibody. Sections were mounted, and cover-slipped. Labeled neurons and their projections were imaged using brightfield and epifluorescence microscopy and a reconstruction of its location as well as of its soma, dendrites and axonal projections was reconstructed using a camera lucida, coupled to a microscope. Axonal trajectories were reconstructed through the dorsal tegmental bundle and ansa peduncularis to the bed nucleus of the stria terminalis (BSTov) and amygdala. Digital micrographs were adjusted for brightness and contrast only.

### Author Contributions

LZ, VSH conceived and designed the study. LZ, VSH, PDSC performed experiments and collected data. LZ secured funding and wrote the original draft. All authors have read and approved the final version of the manuscript.

## Acknowledgements

This research was supported by funding from the following sources: the National Autonomous University of Mexico (UNAM-PAPIIT-IG200121); the Mexican Secretaría de Ciencias, Humanidades, Tecnología e Innovación (Secihti, CF-2023-G-243). We thank Ruud Buijs for discussion and insigtful experimental sugestions, Lee Eiden for reagent support, Mercedes Perusquia-Nava, Arturo Hernández-Cruz and Tatiana Fiordelisio-Coll for equipment supports and Maria Neves Herrera-Mundo and Edgar Jiménez Díaz for technical assistance.

## Notes

### Competing Interest Statement

The authors have declared no competing interest.

### Summary of Updates

We have amended the manuscript with some references and an additional figure (Fig. 5).

